# Germline silencing of UASt depends on the piRNA pathway

**DOI:** 10.1101/290726

**Authors:** Yi-Chun Huang, Henry Moreno, Sarayu Row, Dongyu Jia, Wu-Min Deng

**Author notes:** Department of Biology, Georgia Southern University, Statesboro, GA 30460-8042.

## Abstract

One of the most extensively used techniques in *Drosophila* is the Gal4/UAS binary system, which allows tissue-specific misexpression or knockdown of specific genes of interest. The original UAS vector, UASt, can only be activated for transgene expression in somatic tissues and not in the germline cells. Rørth (1998) generated UASp, a modified UAS vector that is responsive to Gal4 in both somatic and germline tissues, by replacing both the *hsp70* promoter and the SV40 3’UTR with the P transposase promoter and the K10 3’UTR respectively. At present, the mechanisms by which UASt is silenced in germline cells are not fully understood. Here, we report that the piRNA pathway is involved in suppressing UASt expression in ovarian germline cells. Individually knocking down or mutating components of the piRNA biogenesis pathway (e.g., Piwi, AGO3, Aub, Spn-E, and Vasa) resulted in the expression of the UASt-reporter (GFP or RFP) in the germline. An RNA-seq analysis of small RNAs revealed that the *hsp70* promoter of UASt is targeted by piRNAs, and in the *aub* mutant ovary, the amount of piRNAs targeting the *hsp70* promoter is reduced by around 40 folds. In contrast, the SV40 3’UTR of the UASt, which happens to be targeted by the Nonsense-mediated RNA decay (NMD) pathway, is not responsible for germline UASt suppression, as UASt-reporters with NMD-insensitive 3’UTRs fail to show germline expression. Taken together, our studies reveal a crucial role of the piRNA pathway, potentially via the suppression of the *hsp70* promoter, in germline UASt silencing in *Drosophila* ovaries.

## Introduction

The success of the fruit fly *Drosophila melanogaster* as a model organism is heavily attributed to the expansive range and multitude of genetic and molecular tools available to modify gene expression at will. One such commonly used genetic tool is the transgenic Gal4/UAS system, designed for targeted gene expression (Brand and Perrimon, 1993), which allows ectopic expression of any gene (or transgene) in specific tissues, independent of their native regulators. The yeast Gal4 gene was inserted into the *Drosophila* genome under various enhancers and no deleterious effects were observed even when expressed in high levels, making it a ‘safe’ tool to use (Ryder and Russell, 2003). The system relies on the Gal4 gene product binding to and activating the Upstream Activating System (UAS). A gene of interest, inserted downstream of the UAS, will only be expressed if the Gal4 protein is first expressed and then binds to the UAS sequence.

The Gal4/UAS system remains one of the most useful and adaptable tools available; Duffy (2002) called it “a fly geneticist’s Swiss army knife”. However, the original UAS (UASt) is not expressed in the germline cells of the *Drosophila* ovary (Brand and Perrimon, 1993). *Drosophila* ovaries are an extensively used model system for developmental and genetic studies and are ideal for analyzing signaling pathways and complex cellular mechanisms during oogenesis (Velentzas *et al*. 2015). In 1998, Rørth modified various components of the UAS vector, named it ‘pUASp’ and called the original UAS vector ‘pUASt’. UASp has 14 Gal4-binding sites and a GAGA site which allows the target element to transpose efficiently (Rørth, 1996), while UASt has 10 Gal4-binding sites and no GAGA site (see illustration in Rørth, 1998). In order to have germline expression of UASp, the *hsp70* promoter on UASt was replaced by a transposase promoter, since it has high germline expression during oogenesis (Rørth 1998). In addition, the termination sequence source was changed from SV40 3’UTR to K10 to prevent the destabilization of the expressed transcripts (Serano et al., 1994). Given the number of changes made to the original UAS system, it is unclear what is the exact mechanism of silencing in the germline.

Transposable elements (TEs, also called transposons) are mobile selfish DNA elements that exist in the genome of most eukaryotes. TEs take advantage of the host cellular machinery to replicate within the tissue and can result in mutations and chromosome instability (Halic and Moazed, 2009). Host organisms have evolved multiple mechanisms to control the mobilization of TEs to maintain genome integrity. One of these defense systems involves PIWI-interacting RNAs (piRNAs), which function primarily in germline tissues (Siomi et al., 2011). In *Drosophila*, piRNA biogenesis employs a unique ‘ping-pong cycle’ mechanism for piRNA processing and amplification in the *nuage*, a perinuclear structure surrounding the nurse cells in developing egg chambers (Saito et al., 2006; Brennecke et al. 2007; Malone et al. 2009; Siomi et al., 2011). The localization of *nuage* related proteins such as Vasa, Aub, and AGO3 can be used as an indicator for the piRNA biogenesis pathway (Findley et al., 2003; Lim and Kai, 2007; Malone et al., 2009; Handler et al., 2013; Lo et al., 2016).

In this study, we compared each component of the UASt and UASp vectors to determine which elements on the UASt vector are involved in its suppression in the germline. Our findings reveal that the interaction between piRNAs and the *hsp70* promoter is responsible for the suppression of UASt-transgene expression in the germline cells, and the interaction between NMDs and the SV40 3’ UTR is unlikely to have a role in UASt germline suppression.

## Materials and Methods

### Fly stocks and genetics

The following fly stocks were used in this study: *act>CD2>Gal4, UASt-RFPnls* (BL30558); *UASt-mRFP* (BL3417); *UASt-GFP*^*nls*^ (BL4776); *pGW-HA* (a gift from S. Yamamoto, Baylor College of Medicine); *Coin-FLP/Gal4* (BL59268); *UASp-mGFP* (BL58721); *mat-Gal4* (BL7062); *AGO3*^*t2*^ (BL28269); *AGO3*^*t3*^ (BL28270); *aub*^*QC42*^ (BL4968); *aub*^*HN2*^ (BL8517); RNAi lines used in this study are listed in Tables 1 and 2. Flies were maintained and raised at 25°C. Adult female flies for Flp-out experiments were heat-shocked for 30 minutes at 37°C. Two days after heat shock, the flies were dissected to harvest ovaries. The collected ovaries were subjected to immunofluorescence staining.

**Table 1.** Trip lines of targeted genetic screen.

| Category | Genes | Stocks | UAS<sup>t</sup> Expressed in Germline |
| --- | --- | --- | --- |
| **Control & randomly selected** | Luciferase | BL35788 | NO |
|  | psi | BL34825 | NO |
|  | Dl | BL34322 | NO |
| **small RNA** | **su(var)210** | BL29448 | NO |
|  |  | BL31623 | NO |
|  |  | BL32956 | **YES** |
|  |  | BL32915 | **YES** |
|  | su(var)3-9 | BL32914 | NO |
|  | su(var)205 | BL33400 | NO |
|  | **piwi** | BL33724 | **YES** |
|  | **spn-E** | BL34808 | **YES** |
|  | **aub** | BL39026 | **YES** |
|  | tejas | BL41928 | NO |
|  | AGO1 | BL33727 | NO |
|  | **AGO3** | BL44543 | **YES** |
|  | **vasa** | BL34950 | **YES** |
|  | kr | BL34632 | NO |
|  | me31B | BL33675 | NO |
|  | **zuc** | BL36742 | **YES** |
|  | R2D2 | BL34784 | NO |
| **Insulators** | mod(mdg4) | BL32995 | NO |
|  |  | BL33907 | NO |
|  | su(HW) | BL34006 | NO |
|  |  | BL33906 | NO |
|  | CP190 | BL33903 | NO |
|  |  | BL33944 | NO |
|  | CTCF | BL40850 | NO |
|  | BEAF-32 | BL29734 | NO |
|  | GAF/Trl | BL40940 | NO |
| **Polycomb** | E(z) | BL33659 | NO |
|  | Su(z)12 | BL33402 | NO |
|  | Pc | BL33622 | NO |
|  |  | BL33964 | NO |
|  | pea | BL32838 | NO |
| **Hsp protein** | Hsp70B | BL32997 | NO |
|  |  | BL33948 | NO |
|  | Hop | BL34002 | NO |
|  |  | BL32979 | NO |
|  | **Hsp83** | BL33947 | **YES** |

**Table 2.** Genes involved in NMD regulation.

| Category | Genes | Stocks | UAS<sub>t</sub> Expressed in Germline |
| --- | --- | --- | --- |
| **NMD** | Upf1 | BL43144 | NO |
|  | Upf2 | BL31095 | NO |
|  | Upf3 | BL44565 | NO |
|  |  | BL58181 | NO |
|  | Smg1 | BL41945 | NO |
|  |  | BL35349 | NO |
|  | Smg5 | BL31090 | NO |
|  |  | BL62261 | NO |

### Antibodies, immunofluorescence staining and confocal microscopy

Immunocytochemistry was carried out as described previously (Deng *et al.*, 2001). The following antibodies were used: rat anti-Vasa (1:300; Development Studies Hybridoma Bank), rabbit anti-HA-tag (C29F4, 1:100; Cell Signaling). Secondary antibodies were stained with Alexa Fluor® 546 or 488 and nuclear DNA was labeled with DAPI (Invitrogen). Images were acquired with a Zeiss LSM-800 confocal microscope and assembled in Adobe Illustrator.

### Quantitative RT-PCR analysis

Total RNA from two-day-old *Drosophila* ovaries was isolated using Trizol Reagent (Invitrogen) according to the manufacturer’s instructions and then treated with 2 U/µl of DNase I (Ambion) for 30 minutes at 37°C. One microgram of total RNA was reverse-transcribed in 20 µl of reaction mixture containing Superscript II reverse transcriptase (Invitrogen) and oligo (dT)12-18 primer according to the protocol for Superscript II first-strand cDNA synthesis system. One microliter cDNA (reverse transcribed from 50 ng of RNA) was subjected to quantitative real-time PCR (in 25 µl reaction volume) by using primers specific to a transposable element or RP49 (primer sequences below) and cDNA templates were amplified using the Platinum SYBR Green qPCR SuperMix UDG kit, according to the manufacturer’s instructions (Invitrogen). PCR conditions are: 95°C for 10 minutes; 40 cycles of 95°C for 30 seconds, 58°C for 15 seconds, and 68°C for 45 seconds. Real-time PCR was performed using the ABI 7500 Thermocycler (Applied Biosystems), and results were analyzed using SDS version 2.1 software (Austin Biodiversity Web site gallery). Data analysis was done using the 2–ΔΔCT method for relative quantification. Calculated expression values of cDNA samples were normalized to RP49.

#### Primers

Het-A: 5’-ATCCTTCACCGTCATCACCTTCCT-3’, 5’-GGTGCGTTTAGGTGAGTGTGTGTT-3’ and rp49: 5’-ATGACCATCCGCCCAGCATAC-3’, 5’-CTGCATGAGCAGGACCTCCAG-3’ (Pane *et al*., 2007)

RFP: 5’-CATCCCCGACTACATGAAGCTGT-3’, 5’-GCCCTTGAACTTCACCTTGTAGATG-3’

### Small RNA library preparation and analysis

Small RNA libraries were prepared according to instructions for Illumina TrueSeq Small RNA sample prep kit; the library preparation and data analysis were as described in Lo et. al. (2016). Small RNA reads (>22nt) that passed quality control and the removal of rRNAs, snoRNAs and tRNAs reads by bowtie (-a --best --strata -v 1 --un) (bowtie-bio.sourceforge.net/index.shtml) were mapped to the transgene *UASt* nucleotide sequence (detail sequence obtained from Addgene, Cambridge, MA) (-a -v 0 -m 1) by bowtie.

## RESULTS

### UASt-reporter showed germline expression in Su(var)2-10 knockdown

The bipartite Gal4/UAS system is one of the most important and widely used genetic tools in *Drosophila*. It is primarily used for *in vivo* overexpression and misexpression of genes or RNAi mediated gene knockdowns. However, the original UASt construct is not active in the germline cells of the *Drosophila* ovary; as demonstrated here when the *UASt-RFP*^*nls*^ transgene was driven by a ubiquitously expressed Gal4 under the *Actin5C* promoter (*act-Gal4*), RFP expression was only detected in the somatic follicle cells, but not in the germline nurse cells or the oocyte (Fig 1A). The UASp, a modified vector designed by Rørth (1998), by contrast, can be activated in both somatic and germline cells, as shown here the *UASp-mCD8GFP* transgene, driven by the same *act-Gal4*, showed expression in both the follicle cells and nurse cells in the ovary (Fig 1A’).

**Figure 1.**
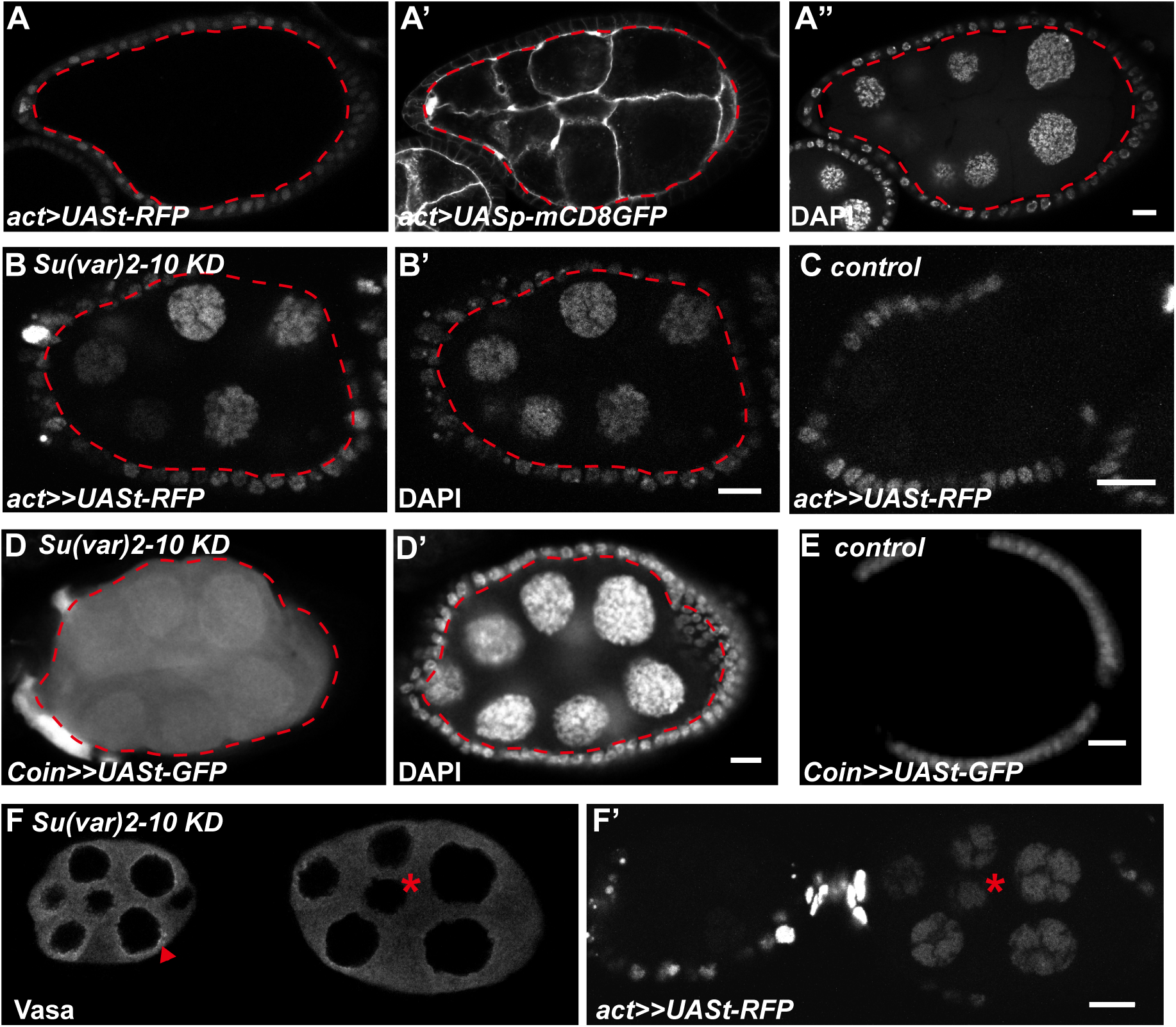
Ectopic germline expression of UASt in *Su(var)2-10* knockdown egg chambers. Broken red lines mark the germline. (A-A”) Gal4 expressed under *act* promoter: UASt-RFP was observed only in follicle/somatic cells; however, UASp-mCD8GFP was expressed in both somatic and germline cells in the same egg chamber. (B, B’) Flp-out Gal4 with *Su(var)2-10* KD: UASt-RFP was now also observed in germline cells. (C) Control Flp-out Gal4: UASt-RFP does not have germline expression. (D, D’) CoinFLP-Gal4 with *Su(var)2-10* KD: ectopic germline UASt-GFP expression; compare to control (E). (F, F’) Wild-type egg chamber (left) and Su9var)2-10 knockdown egg chamber (right) stained for the Vasa protein. Control egg chamber has Vasa localization in the nuage (red arrow head); *Su(var)2-10* KD (indicated by an asterisk, and with germline UASt-RFP expression in F’) had no/reduced Vasa protein in nuage). Nuclei were labeled with DAPI. Posterior is to the right. Scalebars 10 µm.

In a genome-wide *in vivo* RNAi screen to identify genes that potentially regulate Notch signaling in follicle cells (Jia et al., 2015), we used the Flp-out Gal4/UAS system (*act>CD2>Gal4, UASt-RFP*^*nls*^) (Ito *et al*., 1997; Pignoni and Zipursky, 1997) to generate knockdown mosaics in ovarian cells. Although the RFP reporter expression is restricted to the somatic cells since it is cloned on the UASt vector, the collection of TRiP RNAi lines used for this experiment are effective in both the somatic and germline cells (Ni *et al.*, 2011). Unexpectedly, we found that the RFP signal was detected in some nurse cells in mosaic egg chambers with *Su(var)2-10* (CG8086) knockdown (Fig 1B). Consistently, when the CoinFLP system (Bosch et. al., 2015), which also carries the *Actin5C* promoter to generate mosaic Gal4 expression, was used to drive *Su(var)2-10*-TRiP-RNAi expression, UASt-GFP expression was also detected in the nurse cells (Fig 1D). These results suggest that *Su(var)2-10* plays a role in suppressing UASt-transgene expression in the germline.

### The piRNA pathway in germline suppression of UASt

*Su(var)2-10* has been shown to be potentially involved in piRNA biogenesis, chromosome stability and epigenetic regulations (Hari *et al*., 2001; Muerdter *et al*., 2013). We asked if any of these molecular functions are disrupted in the ovarian germline cells when *Su(var)2-10* is knocked down, and found that Vasa localization in the *nuage* in nurse cells (Fig 1F) was significantly reduced. This phenocopies the piRNA pathway mutants (Malone *et al*., 2009), suggesting that *Su(var)2-10* is a part of the piRNA biogenesis machinery.

Next, we performed a candidate RNAi screen of genes involved in small RNA biogenesis, epigenetic regulation, or heat shock responses. This was to determine if any of them would suppress UASt-reporter expression in the germline, in a similar manner to *Su(var)2-10*. Among the 28 genes examined (Table 1), six belonged to the piRNA biogenesis pathway and all of them showed some UASt-RFP expression in germline nurse cells when knocked down using the Flip-out Gal4 to generate mosaics (*piwi* (Fig 2A), *aub* (Fig 2B), *AGO3* (Fig 2C), *spn-E* (Fig 2D), *vas* (Fig 2E), or *zuc* (Fig 2F)). Interestingly, we also found germline expression of UASt-RFP when *Hsp83* was knocked down (Fig 3A, circled with a dashed line). *Hsp83* has been reported to be involved in piRNA biogenesis, and is normally enriched in the *nuage* (Olivieri *et al*., 2012). As expected, we observed Vasa mislocalization when *Hsp83* was knocked-down (Fig 3A’, arrowheads), a phenotype that is consistent with piRNA gene mutations. In contrast, we did not detect any UASt-RFP expression in the germline cells in control egg chambers with Luciferase knockdown (n = 423, Table 1).

**Figure 2.**
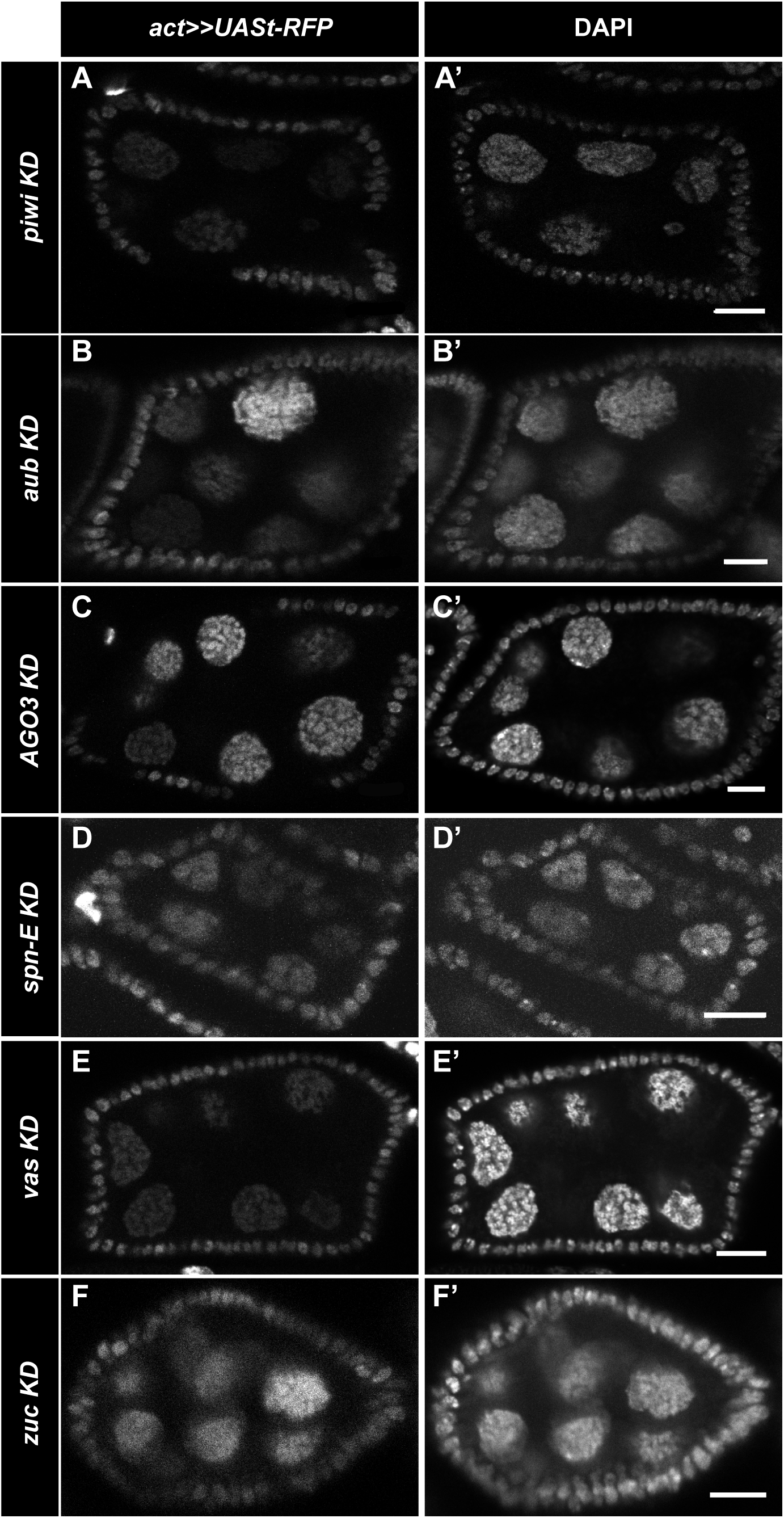
piRNA pathway suppresses germline UASt expression. Targeted genetic screen with Flp-out *act-Gal4* found that UASt-RFP can be expressed in germline cells when KD of following piRNA components (A) piwi, (B) aub, (C) AGO3, (D) spn-E, (E) vas, and (F) zuc. Nuclei were labeled with DAPI. Posterior is to the right. Scalebars 10 µm.

**Figure 3.**
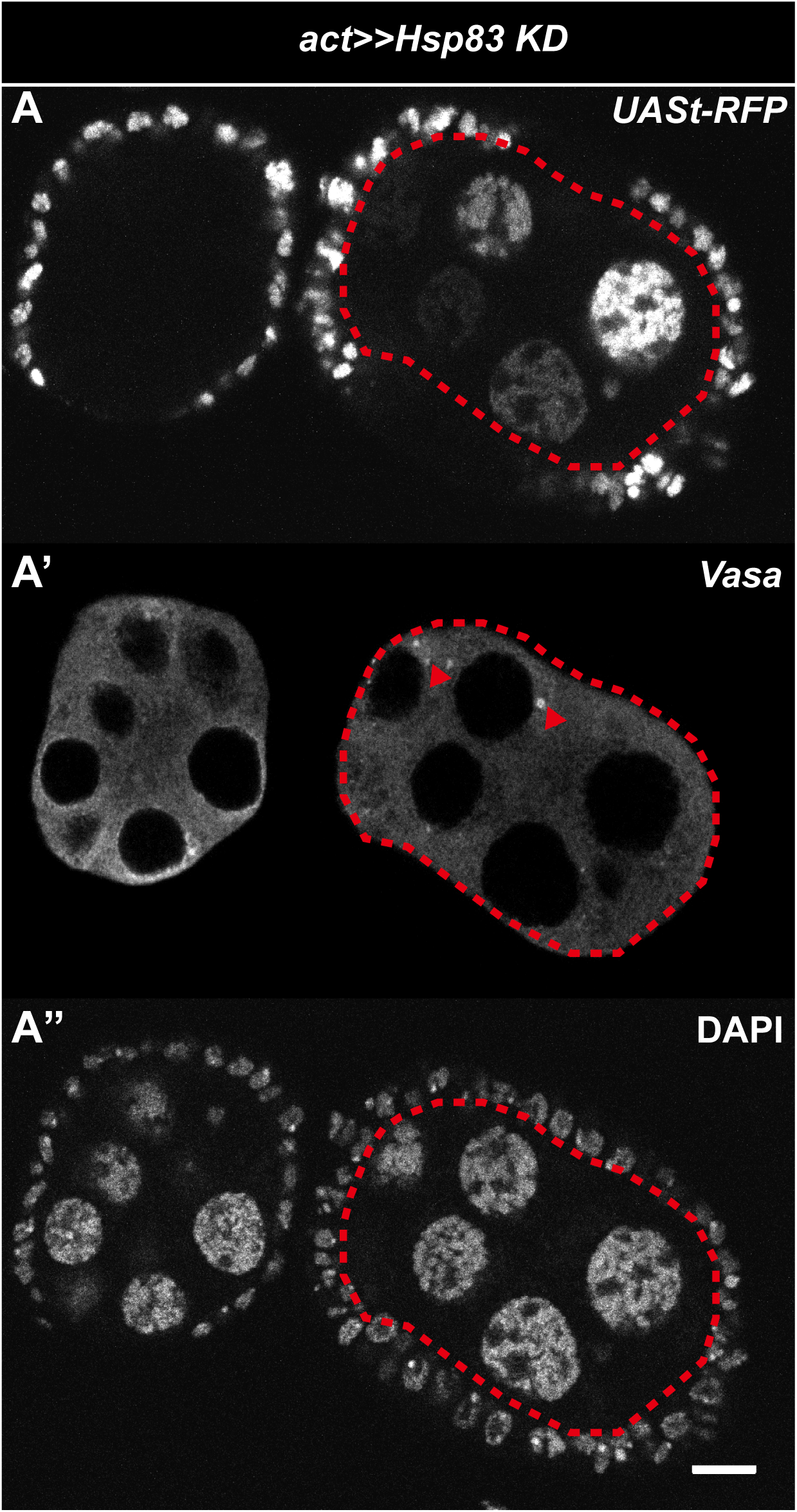
*Hsp83* suppresses germline UASt expression via the piRNA pathway. *Hsp83* Knockdown egg chamber (right); compare to control egg chamber (left). *Hsp83* KD egg chamber has germline *UASt-RFP* expression (A) and mislocalized Vasa protein (arrowhead) (A’) compared to wild-type Vasa localization in (A’). Nuclei were labeled with DAPI. Posterior is to the right. Scalebars 10 µm.

We then examined UASt-reporter expression in the germline of mutants defective in piRNA production. We expressed *act-Gal4* driven UASt-RFP (*act>UASt-RFP*) in a trans-heterozygous mutant of *AGO3* (*AGO3*^*t2/t3*^), and found that all *AGO3*^*t2/t3*^ mutant ovaries expressed RFP in both somatic and germline cells (Fig 4B, 100%, n = 57). By contrast, *AGO3*^*t2/+*^ heterozygous egg chambers showed RFP expression only in the somatic follicle cells (Fig 4A). We also tested the expression of UASt-RFP driven by *mat-tub-Gal4*, a germline specific Gal4, in *aub*^*QC42/HN2*^ trans-heterozygous mutants. As expected, RFP expression was detected in both the nurse cells and the oocyte during oogenesis (Fig 4D). Taken together, these results indicate that the suppression of UASt-transgene expression in the germline depends on the piRNA biogenesis pathway.

**Figure 4.**
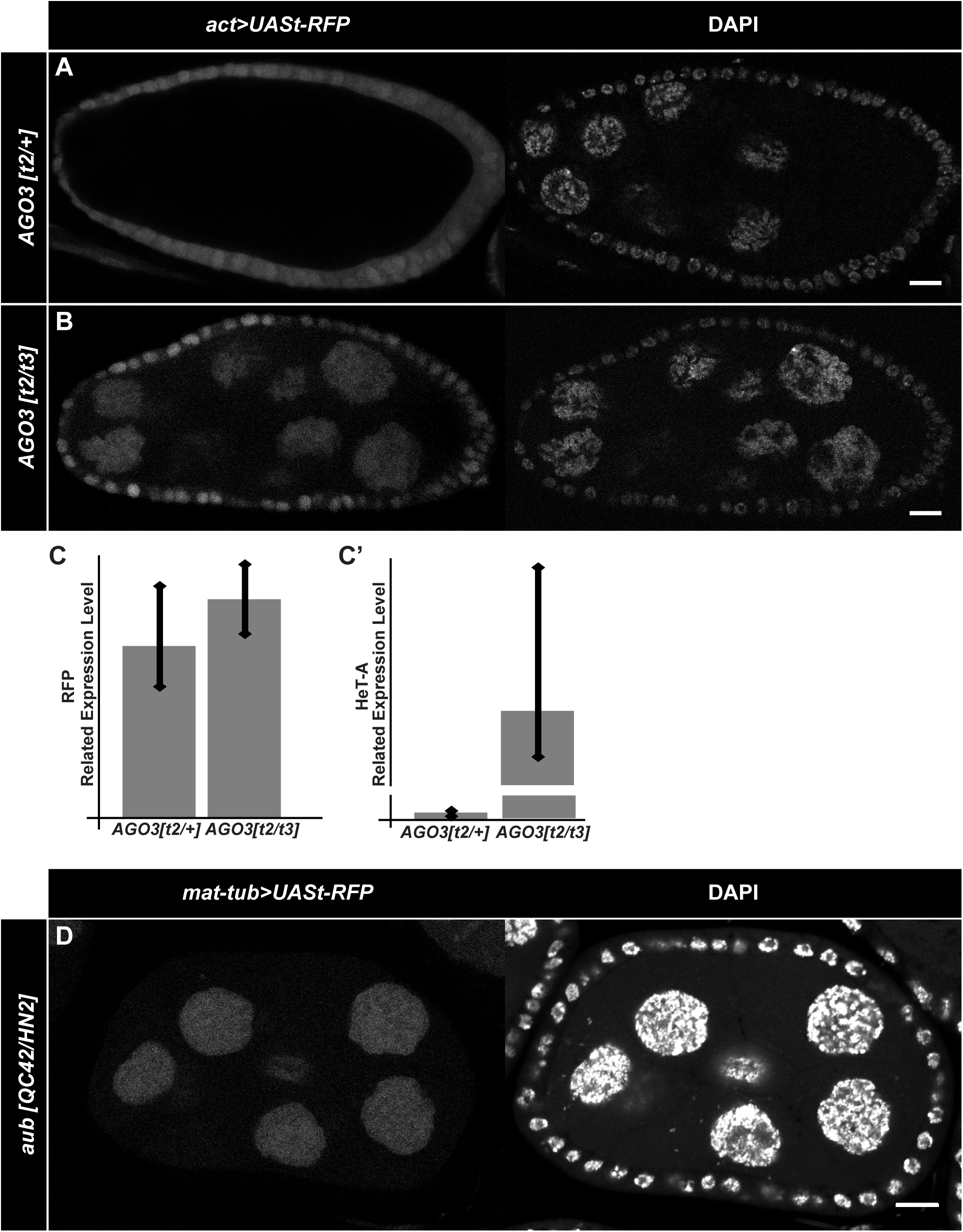
Silencing of germline UASt-transgene expression is transcriptionally regulated. (A) Heterozygous *AGO3*^*t2*^ allele showed only follicle/somatic expression of UASt-RFP driven by *act-Gal4*. (B) Trans-heterozygous *AGO3*^*t2/t3*^ egg chambers displayed both somatic and germline UASt-RFP expression. (C) qRT-PCR results. Overall RNA level of RFP from trans-heterozygous *AGO3*^*t2/t3*^ ovaries was higher (1.3 fold), compared to heterozygous *AGO3*^*t2*^ ovaries. Increased HeT-A expression (60 fold) indicated the piRNA pathway was inactive in *AGO3*^*t2/t3*^ mutant alleles. (D) UASt-RFP driven by *mat-tub-Gal4*, a germline specific driver, also displayed germline UASt-RFP expression in trans-heterozygous *aub*^*QC42/HN2*^ egg chambers. Nuclei were labeled with DAPI. Posterior is to the right. Scalebars 10 µm.

To determine whether UASt-RFP suppression is at the transcription level, we performed quantitative RT-PCR (qRT-PCR) analyses, and found the RFP transcript level was 1.3-fold higher in *AGO3*^*t2/t3*^ trans-heterozygous ovaries than the *AGO3*^*t2/+*^heterozygous controls. As a control, we examined the expression of Het-A, a transposable element that is targeted by piRNAs (Brennecke *et al*., 2007), and found a 60-fold increase of the Het-A transcript in *AGO3*^*t2/t3*^ mutant ovaries (Fig 4C’), indicating that piRNA production is indeed strongly suppressed in these ovaries. Together, these results suggest that germline expression of UASt-RFP is suppressed by piRNAs at the transcriptional level (Fig 4C).

### The hsp70 promoter is a piRNA target

To determine exactly how piRNAs suppress UASt-transgene expression in the germline, we performed a RNA-seq analysis of small RNAs from ovaries of *aub*^*QC42/HN2*^ and *w*^*1118*^ flies. After mapping the piRNAs from the RNA-seq analysis on to the pUASt sequence, we found that the *hsp70* promoter was heavily targeted by piRNAs in *w*^*1118*^ flies (Fig 5A, B). This result is consistent with the report that the *hsp70* locus itself provides the substrates for high piRNA production in transgenic lines (Olonikov *et al.*, 2013). By contrast, *aub*^*QC42/HN2*^ ovaries had a significantly reduced level (∼40-fold lower) of piRNAs targeting the *hsp70* promoter sequence compared with the *w*^*1118*^ controls (Fig. 5B). These findings, along with the qRT-PCR results that *AGO3*^*t2/t3*^ mutant ovaries had elevated levels of the UASt-RFP transcript, suggest that the *hsp70* promoter in the UASt vector is targeted by piRNAs from the *hsp70* locus, thus suppressing UASt-transgene transcription in the germline cells.

**Figure 5.**
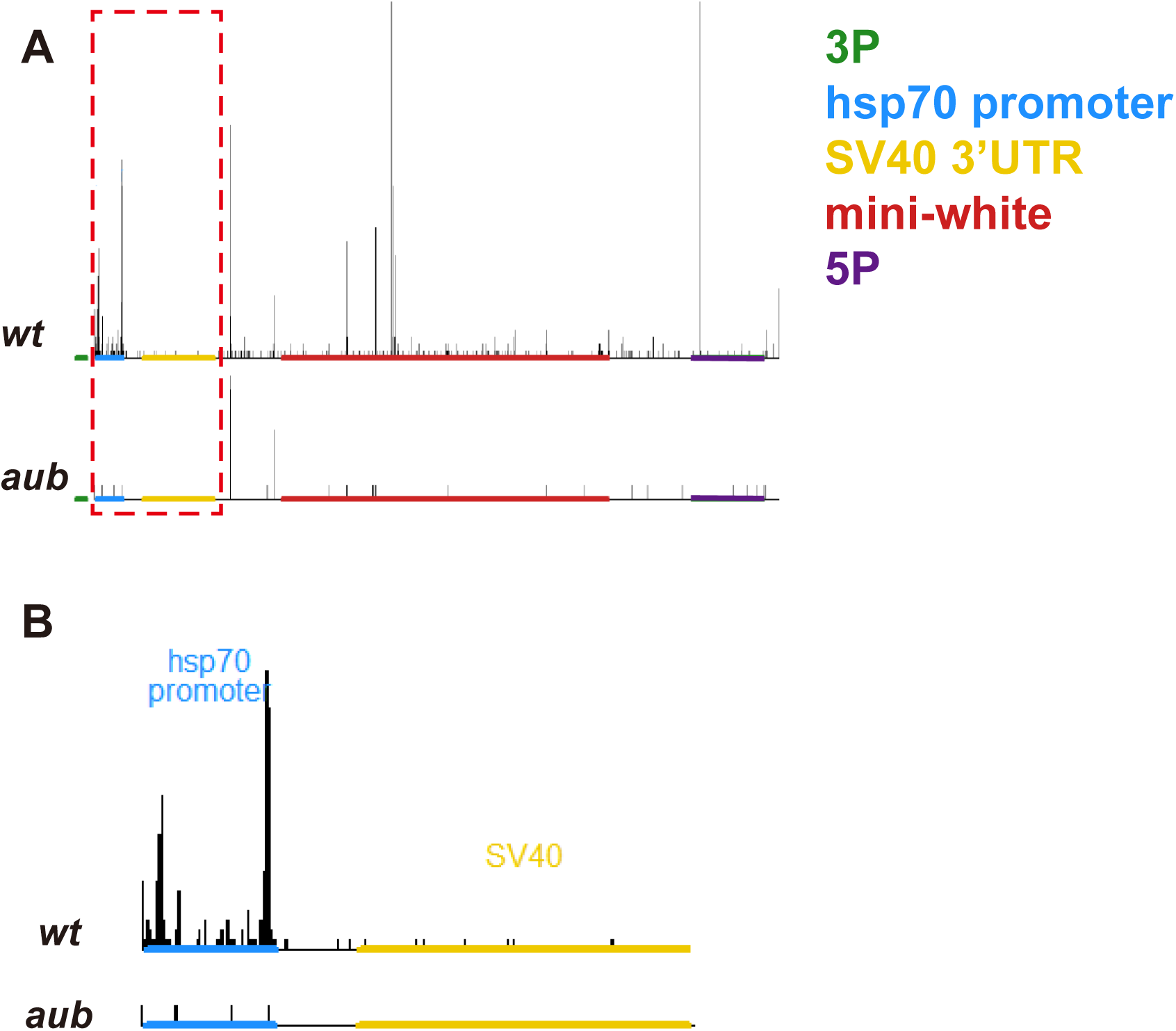
hsp70 promoter is a piRNA target. Small RNA deep sequencing from wild-type and *aub*^*QC42/HN2*^ ovaries were mapped to the UASt sequence. X axis: position of the major compositions of the UASt. Y axis: normalized aligned read counts for piRNA. (A) The piRNA sequencing reads (>22 nt) from ovary samples for wild-type (top plot) and trans-heterozygous *aub*^*QC42/HN2*^ (bottom plot). (B) Closed view of *hsp70* promoter and SV40 3’UTR sequences from the mapping result. This figure was generated by R.

### The 3’UTR is not the cause of UASt germline suppression

Besides the promoter, the other major difference between UASt and UASp is the 3’UTR tail. UASt has an SV40 3’UTR, whereas UASp carries a K10 tail (Rørth, 1998). Metzstein and Krasnow (2006) discovered that the nonsense-mediated mRNA decay factors (NMD) targeted the SV40 3’UTR sequence, thereby suppressing the expression of upstream genes. To determine whether NMD is involved in germline UASt suppression, we examined the expression of a modified UASt–GFP, which contains a shortened SV40 3’ UTR excluding the NMD binding site (Metzstein and Krasnow 2006). When driven by *act-Gal4*, this NMD-non-sensitive line (K45) showed no ovarian germline expression of UASt-GFP (Fig 6A-B). We further tested another modified UASt vector, the pGW construct (Bischof *et al*., 2013), which has *tubulin* 3’ UTR instead of the SV40 3’UTR. Using the Flp-out Gal4 system, we co-expressed UASt-RFP and GW-HA in the ovary, and detected neither RFP nor HA proteins in germline cells (Fig 6C). Additionally, we knocked down genes involved in the NMD pathway (*Upf1, Upf2, Upf3, Smg1*, and *Smg5)*, individually, and no germline UASt-reporter expression was detected in the ovary (Table 2). Taken together, these results suggest that the SV40 3’ UTR and NMD pathway are not involved in germline UASt silencing.

**Figure 6.**
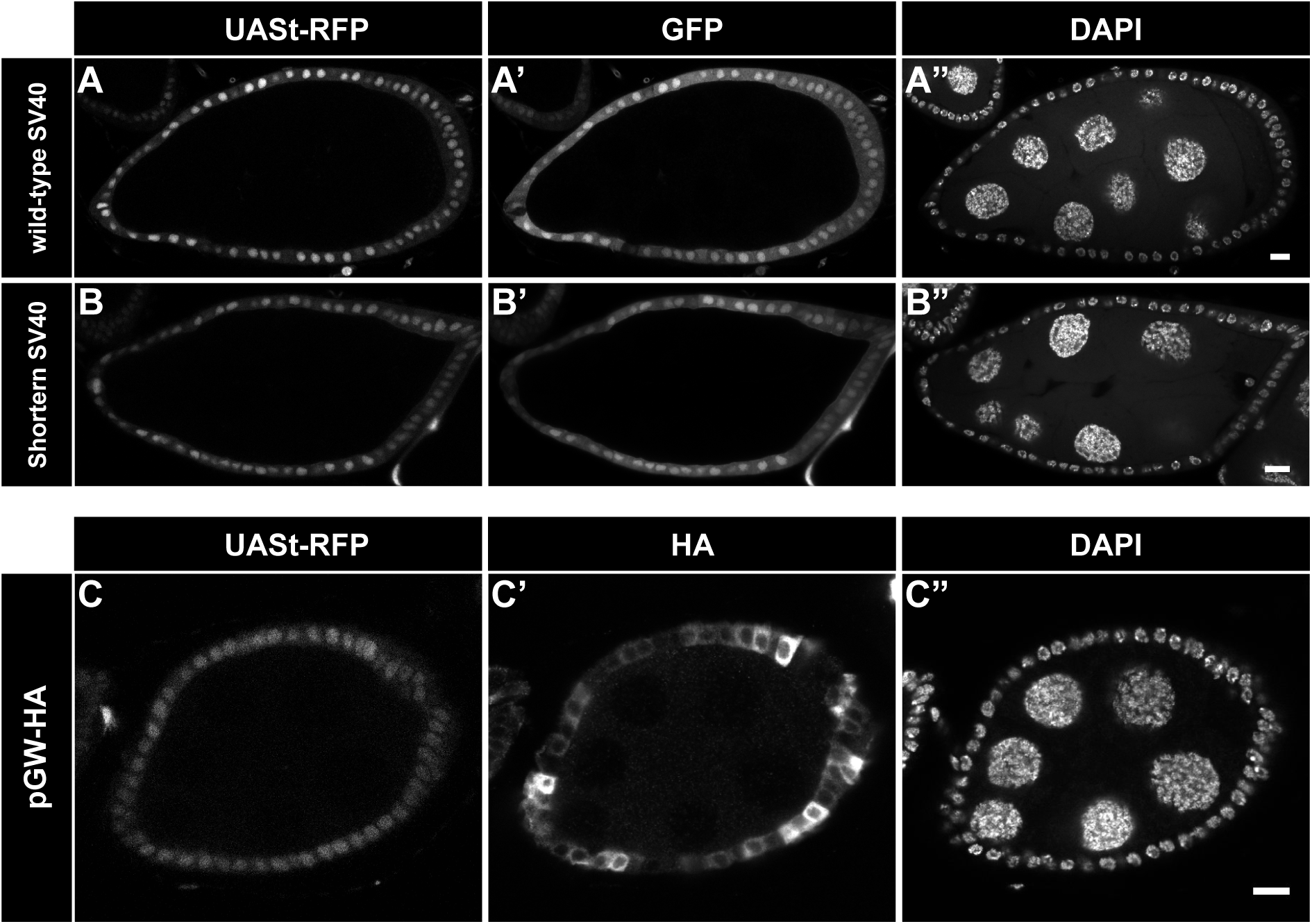
3’UTR of UASt is not involved in germline silencing of UASt. (A) UASt-K37-GFP reporter line, with the wild-type SV40 3’UTR (NMD sensitive), (B) UASt-K45-GFP reporter line, with a shortened SV40 3’UTR (NMD insensitive), and (C) pGW-HA-tag line with a tubulin 3’UTR. (A-C”) Driven by *act-Gal4* along with UASt-RFP. None of the above showed expression of UASt in germline cells. Nuclei were labeled with DAPI. Posterior is to the right. Scalebars 10 µm.

## Discussion

The Gal4/UAS system has been used extensively in a variety of forms and adaptations for a wide range of applications. However, the one downside is that the original UASt is not expressed by germline cells of *Drosophila* ovaries, which limits the experimental power of germline cells. Rørth (1998) succeeded in making a functional ‘pUASp’ system for use in both germline and somatic cells by modifying two important components of the system, the *hsp70* promoter and the SV40 3’ UTR. In this study, we analyzed each component on the UASt that is different from those on the UASp. We show that the *hsp70* promoter targeted by piRNAs is crucial in the silencing of UASt-transgene expression in germline cells. Although the SV40 3’ UTR can be silenced by nonsense-mediated mRNA decay factors (NMD), it did not seem to affect UASt germline expression. From a technical standpoint, this new understanding of how UASt-transgene is silenced in *Drosophila* germline cells could lead to new and better design of transgenes to be expressed in different tissues in this powerful genetic model organism.

piRNAs are abundant in the germline and can repress transposable elements (TE), which prevents the disruption of the genome and reduces the rates of mutation (Ku and Lin, 2014). It has been previously reported that transgene-derived piwi-interacting RNAs (piRNAs) are complementary to the *hsp70* promoter and can cleave and process non-homologous regions of the endogenous *hsp70* transcripts into more piRNAs (Olovnikov *et al.* 2013). Our RNA-seq analysis confirmed that the basic *hsp70* promoter on pUASt is a target for piRNAs, and these piRNAs are significantly reduced when the piRNA biogenesis pathway is disrupted. Hsp70 is the principal inducible heat shock protein in *Drosophila*, with both protective and deleterious roles during development (Feder and Krebs 1998, Zatsepina *et al*. 2001). A previous study proposed the possible involvement of Hsp70 in the biogenesis of piRNA in *Drosophila* following severe heat-shock conditions, but could not conclusively determine the details of this regulatory process (Funikov *et al* 2015). This, in combination with our findings, suggest that piRNAs may be involved in the heat-stress induced response mediated by increased Hsp70 levels for normal development. Future studies will need to address the detailed mechanisms by which piRNAs are able to regulate *hsp70* in this tissue. The overarching question regarding the developmental significance of this differential pattern of *hsp70* expression in the germline and somatic cells of the *Drosophila* ovary remains to be answered.

## Acknowledgements

We thank members of the Deng laboratory for technical support and discussions. We thank S. Yamamoto for suggestions and fly stocks; M. Metzstein, the Developmental Studies Hybridoma Bank, and Bloomington *Drosophila* Stock Center (BDSC) for antibodies, and fly stocks. We thank the Biology Imaging Laboratory, the Molecular Core Facility, the Translational Science Laboratory and the Center for Genomics and Personalized Medicine at Florida State University. We also thank D. Corcoran, P. Michael Albert II and Righting Your Writing, LLC for assistance with editing the manuscript. W.-M. D. is supported by NIH Grant R01GM072562 and NSF Grant IOS-1557904.

